# Inhibition of Stearoyl-CoA desaturase suppresses follicular help T and germinal center B cell responses

**DOI:** 10.1101/601021

**Authors:** Young Min Son, In Su Cheon, Nick P. Goplen, Alexander L. Dent, Jie Sun

## Abstract

Stearoyl-CoA desaturases (SCD) are endoplasmic reticulum (ER) associated enzymes that catalyze the synthesis of the monounsaturated fatty acids (MUFAs). As such, SCD play important roles in maintaining the intracellular balance between saturated fatty acid (SFAs) and MUFAs. The roles of SCD in CD4^+^ T helper cell responses are currently unexplored. Here, we have found that murine and human follicular helper T (T_FH_) cells express higher levels of SCD1 compared to non-T_FH_ cells. Further, the expression of SCD1 in T_FH_ cells is dependent on the T_FH_ lineage-specification transcription factor BCL6. We found that the inhibition of SCD1 impaired T_FH_ cell maintenance and shifted the balance between T_FH_ and follicular regulatory T (T_FR_) cells in the spleen. Consequently, SCD1 inhibition dampened germinal center B cell responses following influenza immunization. Mechanistically, we found that SCD inhibition led to increased ER stress and enhanced T_FH_ cell apoptosis *in vitro* and *in vivo*. These results reveal a possible link between fatty acid metabolism and cellular and humoral responses induced by immunization or potentially, autoimmunity.

## Introduction

Follicular helper T (T_FH_) cells are specialized effector CD4 T helper cells that promote the formation of germinal centers (GCs) and the production of class-switched high-affinity immunoglobulins (1). T_FH_ cells are required for the generation of antibody-mediated protection against microbes, but aberrant T_FH_ responses may cause autoimmunity (2). Therefore, it is important to understand the positive and/or negative regulators of T_FH_ cell responses for the proper induction during vaccination or the inhibition of T_FH_ responses during autoimmunity.

T_FH_ differentiation *in vivo* is a multifactorial multistep process. The transcription factor BCL-6 is considered the master regulator for the differentiation of T_FH_ cells and subsequent germinal center responses (3,4). Interestingly, recent evidence suggests that cell metabolic processes may play important roles in modulating T_FH_ development *in vivo*. BCL6 was shown to repress glycolysis promoting BCL6 expression in activated CD4 T cells (5,6). Consistent with those observations, it has been demonstrated that T_FH_ cells exhibit diminished glycolysis and mitochondrial respiration compared to Th1 cells. However, a recent study has also suggested mTOR kinase complex 1 (mTORC1) related glucose metabolism is required for the T_FH_ generation *in vivo* (7). Therefore, the roles of glucose metabolism in negative and positive regulation of T_FH_ differentiation and/or maintenance remain to be elucidated.

Besides glucose metabolism, emerging evidence has suggested that lipid metabolism also plays important roles in regulating T helper cell responses. The inhibition of acetyl-CoA carboxylase 1 (ACC1), a key substrate for fatty acid synthesis, reduces human and mouse Th17 cell development in favor of regulatory T (Treg) cell differentiation (8). Simvastatin, the inhibitors for 3-hydroxy-3-methyglutaryl (HMG)-CoA reductase, has been reported to promote Treg differentiation whereas it suppresses Th17 development (9). Furthermore, the suppression of fatty acid synthesis (FAS) during T cell activation reduces the development of CD4^+^ T cell memory (10). Currently, the roles of lipid metabolism in T_FH_ differentiation have not been explored.

Stearoyl-CoA desaturase (SCD) is an endoplasmic reticulum (ER) protein that catalyzes the synthesis of monounsaturated fatty acids (MUFAs) from saturated fatty acids (SFAs). Besides the needs of MUFAs for the synthesis of complex lipids such as diacyglycerols (DG), triglycerides (TGs) and cholesterol esters, MUFAs also have important functions in cell signaling and membrane fluidity (11). Therefore, SCD is a highly regulated and conserved enzyme that plays important roles in regulating obesity, insulin resistance, inflammation and cancer progression (12,13). Humans have two SCD isoforms (SCD1 and SCD5), while mouse has four (SCD1-4) that share homology with human SCD1, but not human SCD5 (14,15). The roles of SCD in regulating adaptive lymphocyte responses have not been explored.

Given the importance of lipid synthesis and metabolism to adaptive immunity, we examined the roles of SCD in T_FH_ responses. We found that SCD1 is highly expressed in both human and murine CD4^+^CD44^+^PD1^hi^CXCR5^hi^ T_FH_ cells compared to non-T_FH_ cells. BCL6-deficient CD4^+^ T cells exhibited diminished *Scd1* gene expression. We showed that *in vivo* inhibition of SCD1 activity impaired the maintenance of GC B cells as well as T_FH_ cells. Finally, SCD1 inhibitor increased apoptosis in T_FH_ cells *in vivo* and *in vitro*. Furthermore, inhibition of SCD1 promoted the expression of ER stress genes whereas SCD1 overexpression showed opposite results in T_FH_ cells. Our results suggest that BCL6-mediated SCD1 expression is required for T_FH_ cell maintenance and efficient GC B cell responses *in vivo*.

## Results

### SCD inhibitor suppresses T_FH_ and GC B cell responses

To explore the function of lipid metabolism in regulating T_FH_ and GC B cell responses, we immunized influenza X31 virus and then treated with mice with Statin (an inhibitor of HMG-CoA reductase that is required cholesterol synthesis (16), C75 (an inhibitor of FASN that is required for generation of long-chain fatty acids (17) or A939572 (an inhibitor of SCD1) (18). We checked T_FH_ (CD4^+^CD44^+^PD1^+^CXCR5^+^) and GC B (B220^+^FAS^+^GL-7^+^) cell responses in the spleen at 14 days post immunization. We found that statin and C75 treatment diminished T_FH_, but not GC B cell number following influenza immunization (Fig. 1A). However, SCD inhibitor significantly diminished both the frequencies and the cell numbers of T_FH_ and GC B cells (Fig. 1A, B). SCD inhibitor did not alter the total number of CD4^+^ T and B cells (Fig. 1B), nor the production of Th1 cytokine interferon (IFN)-gamma in CD4^+^ and CD8^+^ T cells (Fig. 1C). These results suggest that SCD inhibition selectively impair T_FH_ but not Th1 cell formation following influenza immunization.

**Figure 1.**
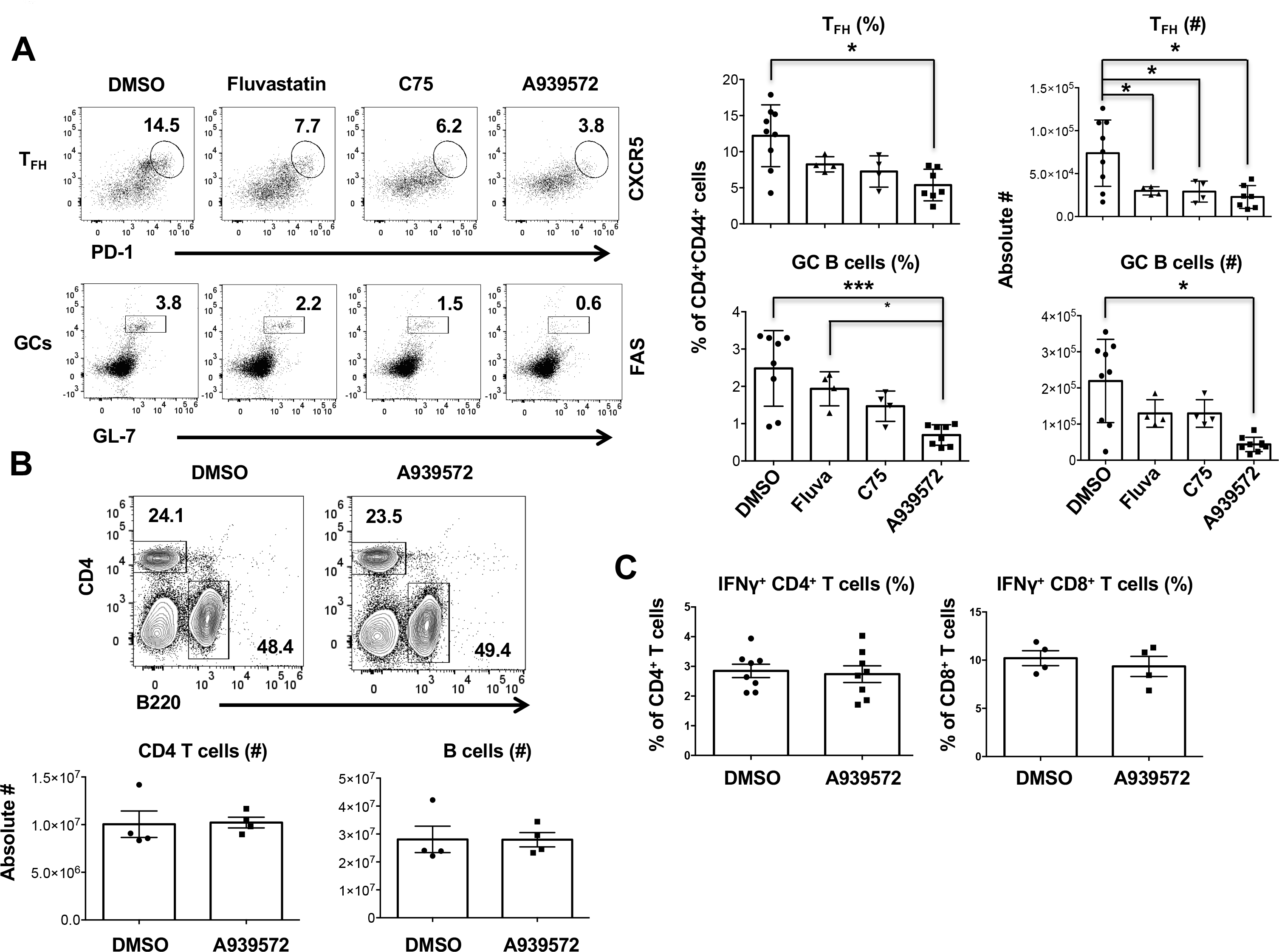
SCD inhibition suppresses T_FH_ but not Th1 responses. WT B6 mice were immunized with X31 and treated with indicated inhibitors start at 3 days post immunization (d.p.i.). **A.** Frequencies and absolute cell number of splenic T_FH_, GC B cells were measured by flow cytometry at 14 d.p.i. (n = 4 - 9). **B.** Frequencies and absolute cell number of splenic CD4^+^ T or B cells at 14 d.p.i. (n = 4 - 8). **C.** IFN-γ production by CD4^+^ or CD8^+^ T cells were measured through intracellular staining (ICS) following re-stimulation with PMA and Ionomycin in vitro through flow cytometry at 14 d.p.i. (n = 4 - 8). Representative data are obtained from at 2 independent experiments. Combine results are obtained from at 2 independent experiments. * indicates significant differences (*P*<0.05), ** indicates *P*<0.01 and *** indicates *P*<0.001.

### *Scd1* is highly expressed by human and murine T_FH_ cells

To identify the expression of *Scd1* in the T_FH_ and non-T_FH_ cells, we first analyzed a publicized microarray data set (GEO# GSE50391) of PD1^+^CXCR5^high^, PD1^+^CXCR5^int^ and PD1^+^CXCR5^−^ from human tonsil (19). We found that PD1^+^CXCR5^high^ T_FH_ cells exhibit significantly higher *Scd1* expression compared to non-T_FH_ cells (Fig. 2A). In another microarray data set (GEO# GSE40068), from sorted lymph node cells of mice immunized with KLH in CFA (20), the mouse T_FH_ cells also had enhanced *Scd* genes expressed relative to non T_FH_ cells. However, the expression of *Fasn* was comparable between Non-T_FH_ and T_FH_ cells (Fig. 2B). We then sorted murine T_FH_ and non-T_FH_ (CD4^+^CD44^+^PD1^−^CXCR5^−^) from spleens of day 14 post influenza (X31) immunized mice. We then examined *Scd1*, *Scd2* and *Fasn* expression in T_FH_ and non-T_FH_ by quantitative RT-PCR. We found that T_FH_ cells exhibited drastic enhanced levels of *Scd1* and modest increased *Scd2* expression compared with non-T_FH_ cells (Fig. 2C). Together, these results reveal that both murine and human T_FH_ cells have enhanced *Scd1* expression.

**Figure 2.**
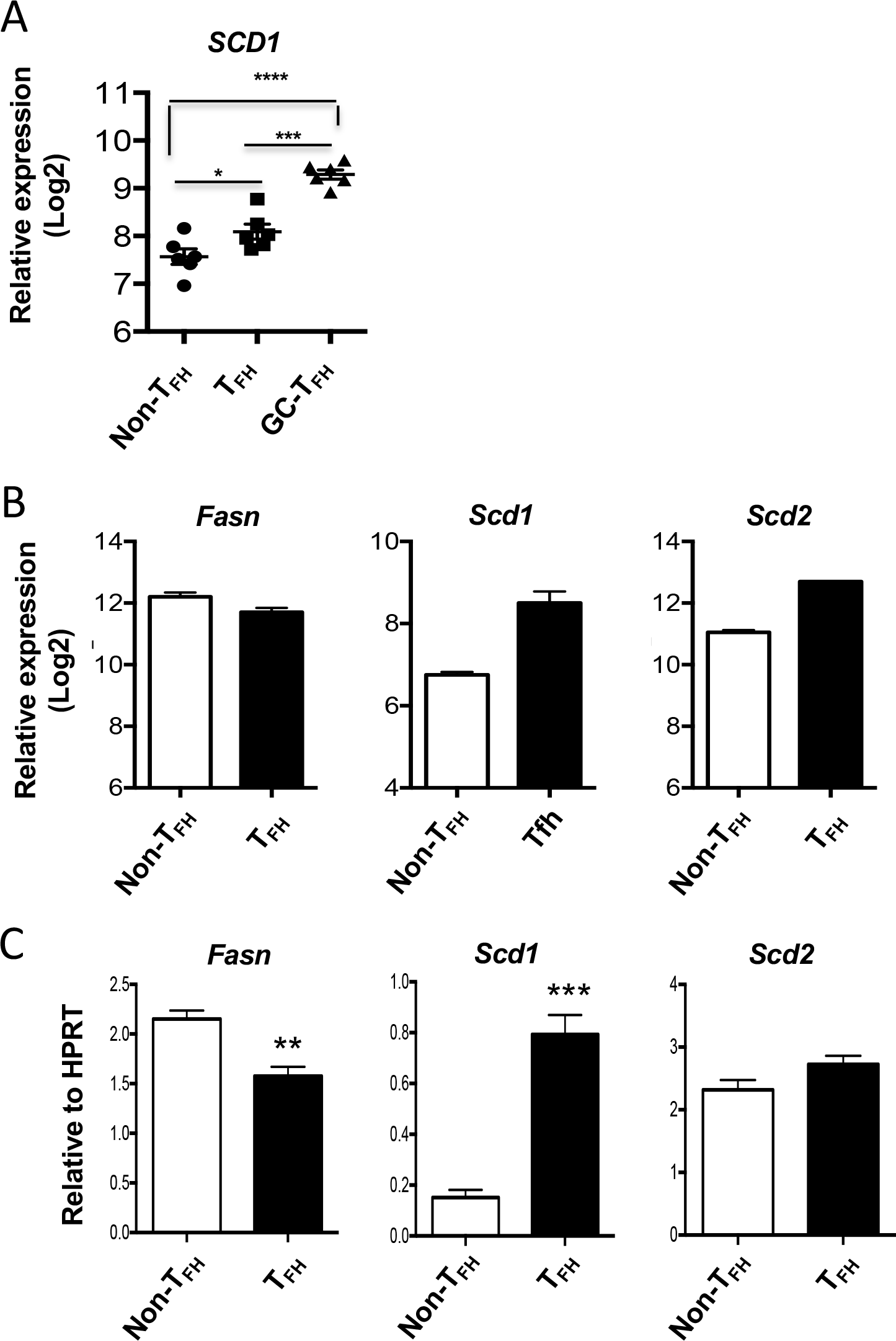
Human and mouse T_FH_ express higher levels of SCD1. (A) Relative SCD1 expression in published microarray data (GEO# GSE50391) CD45RO^+^CXCR5^−^ (Non-T_FH_), CD45RO^+^CXCR5^int^ (T_FH_), and CD45RO^+^CXCR5^hi^ (GC T_FH_) cells from human tonsils. (B) Relative expression of *Fasn*, *Scd1* and *Scd2* in published microarray data (GEO# GSE40068) from BCL6^hi^CXCR5^+^ (T_FH)_ and BCL6^−^CXCR5^−^ (non-T_FH_) cells of the mice immunized with KLH in CFA. (C) Relative expression of *Fasn*, *Scd1* and *Scd2* in sorted splenic T_FH_ (PD-1^+^CXCR5^+^) and non-T_FH_ (PD1^−^CXCR5^−^) cells isolated from X31 immunized mice at 14 d.p.i. Representative results of (C) obtained from at 2 independent experiments. * indicates significant differences (*P*<0.05), *** indicates *P*<0.001 and **** indicates *P*<0.0001.

### BCL6 promotes *Scd1* expression in T_FH_ cells

To determine whether T_FH_ master transcription factor BCL6 is involved in *Scd* expression, we measured *Scd1* and *Scd2* expression in *in vitro* cultured WT or BCL6-deficient (BCL6 KO) CD4^+^ T cells under the T_FH_-like polarization conditions. Both *Scd1* and *Scd2* were diminished in BCL6 KO CD4^+^ T cells (Fig. 3A). We next pre-activated naive CD4^+^ T cells for 2 days and then transduced the cells with BCL6-expressing retroviruses. *Scd1* and *Scd2* were measured in the isolated transduced cells. The BCL6-overexpressed cells exhibited significantly enhanced *Scd1*, and modestly upregulated *Scd2* expression (Fig. 3B). To probe the relationship between BCL6 and SCD1 *in vivo*, we adoptively transferred WT OTII TCR transgenic T cells or BCL6 KO-OTII cells into CD45.1 congenic mice. Following adoptive transfer, mice were immunized with X31-OTII virus. At day 14 post immunization, OTII cells were sorted for measuring *Scd1* and *Scd2* expression. Similar to *in vitro* culture results, the expression of *Scd1* was impaired in BCL6 KO-OTII cells (Fig. 3C). Together, these results demonstrated that BCL6 promotes the expression of *Scd1* in T_FH_ cells.

**Figure 3.**
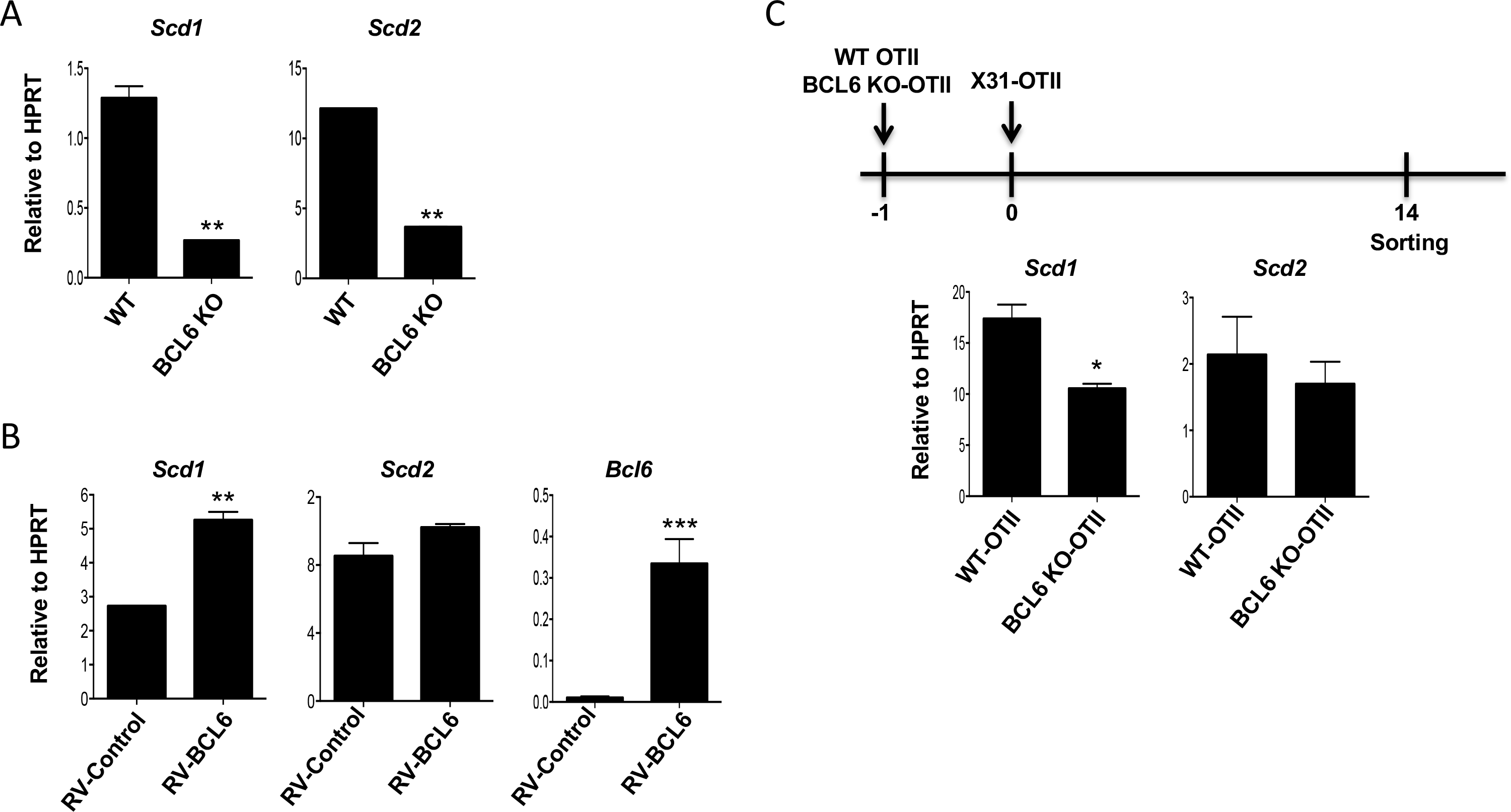
BCL6 promotes SCD1 expression. (A) Relative expression of *Scd1* and *Scd2* in WT or BCL6 KO CD4^+^ T cells under the T_FH_ cell polarizing condition for 4 days. (B) *Scd1*, *Scd2* and *Bcl6* expression in the BCL6-retroviral transduced CD4^+^ T cells under the T_FH_ cell polarizing condition. (C) WT or BCL6 KO-OTII cells were transfer into CD45.1 congenic mice at one day before X31-OTII virus immunization. After day 14 post immunization, the OTII cells were sorted and then *Scd1* and *Scd2* were measured by quantitative RT-PCR (n = 3/group). Representative graphs are obtained from at least 2 (A, C) or 3 (B) independent experiments. Unpaired Student’s t-test: \**P*<0.05; \*\**P*<0.01; and \*\*\**P*<0.001.

### SCD inhibitor suppresses the maintenance of T_FH_ and GC B

Next, we examined whether SCD inhibition is required for development and/or maintenance of T_FH_ and/or GC B cells. To this end, the frequencies and cell numbers of T_FH_ and GC B cells were measured day 8 and day 14 post X31 immunization following SCD inhibition. At day 14 post immunization, SCD inhibitor significantly diminished the number of T_FH_ and GC B cells whereas SCD inhibitor had no effect on T_FH_ and GC B cells at day 8 post immunization (Fig. 4 A and B). The expression of BCL6 and ICOS in the T_FH_ cells between vehicle (DMSO) and SCD inhibitor treated group were comparable (Supplementary Fig. 1). These data suggested that SCD inhibition may not inhibit the formation of T_FH_ and GC B cells, but suppresses the maintenance T_FH_ and GC B cell responses. Recently, T follicular regulatory (T_FR_) cells have been reported as a major regulator repressing excessive T_FH_ and GC B cell responses (21). We found that the frequencies of CD4^+^CD44^+^PD1^+^CXCR5^+^Foxp3^+^ T_FR_ cells were inflated 2-fold relative to T_FH_ cells following SCD inhibition (Fig. 4C). This indicates that SCD inhibition alters the balance of T_FH_ and T_FR_ responses. Consequently, influenza-specific IgG antibody production was impaired in the serum following SCD1 inhibition (Fig. 4D). Together, these results suggest SCD is required for maintenance of T_FH_ and GC B cells through regulation of balance between T_FH_ and T_FR_ cells in the spleen.

**Figure 4.**
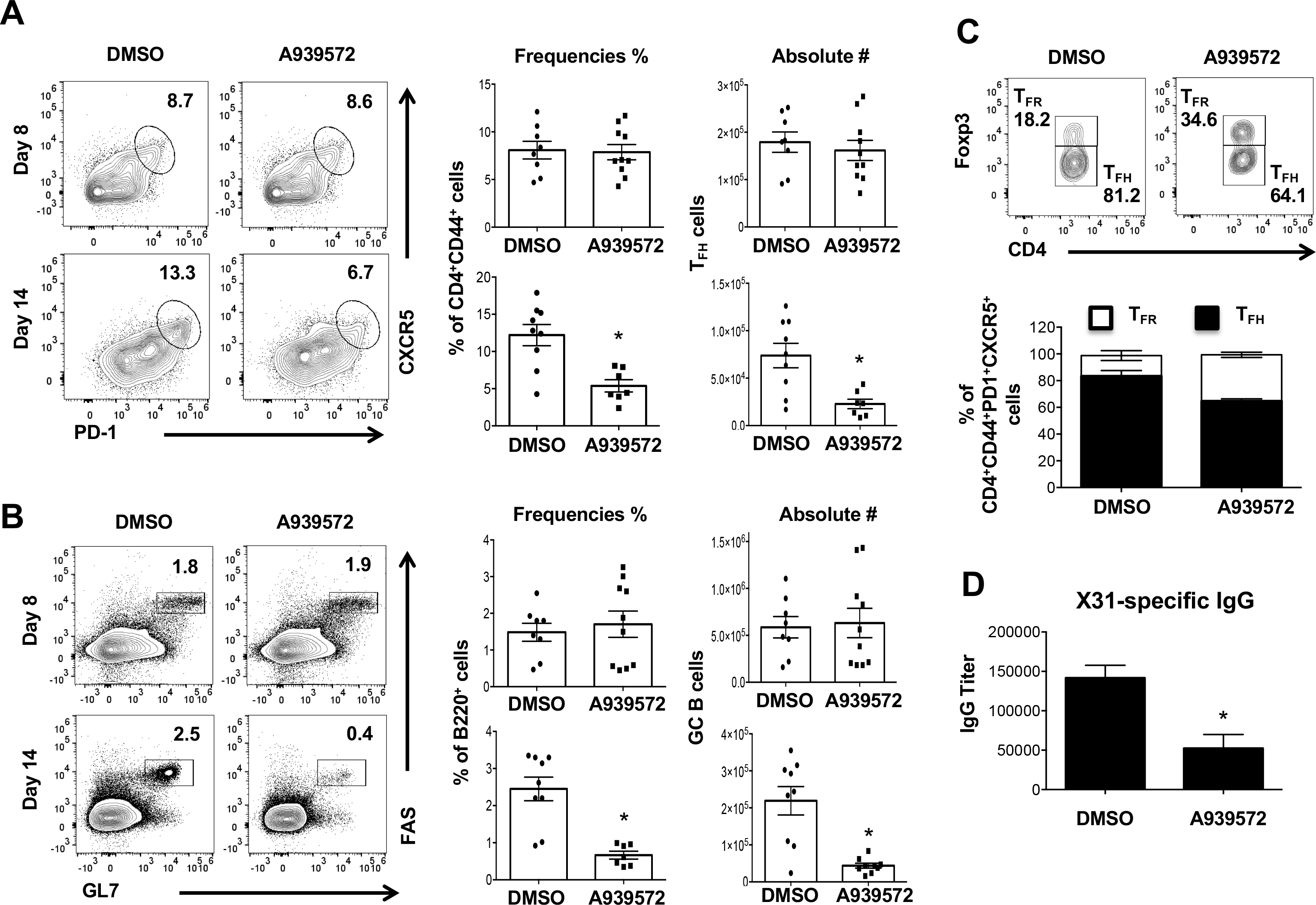
Inhibition of SCD suppresses T_FH_ and GC B maintenance, but promotes T_FR_ responses. (A-B) Frequencies and absolute cell number of splenic T_FH_ and GC B cells in the mice immunized with X31 virus and treated with SCD1 inhibitor at indicated d.p.i. (n = 7 - 10). (C) Frequency of splenic T_FR_ and T_FH_ cells were measured in the in the mice immunized with X31 virus and treated with SCD1 inhibitor at 14 d.p.i. (D) The titer of influenza-specific IgG was measured by ELISA from blood serum at 14 d.p.i. Representative data are obtained from at 3 independent experiments. * indicates significant differences (*P*<0.05) between DMSO control group and SCD inhibitor (A939572) treated group.

### Inhibition of SCD promotes apoptosis and ER stress gene expression

To probe the underlying mechanism, we examined T cell apoptosis following SCD inhibitor treatment. At day 8 post X31 immunization, active caspase 3/7 activities in T_FH_ cells following SCD1 inhibitor treatment were similar to vehicle (DMSO) treated group. However, at day 14 post immunization, caspase 3/7 activities were significantly higher in T_FH_ cells following SCD inhibition compared to DMSO-treated group (Fig. 5A). *In vitro* cell apoptosis was significantly enhanced following SCD inhibitor treatment in a dose dependent manner under Tfh polarizing conditions (Fig. 5B). Taken together, these data demonstrate that SCD activity promotes T_FH_ cell viability. A recent study demonstrated excessive accumulation of SFAs following SCD inhibition promotes endoplasmic reticulum (ER) stress-related cell apoptosis (22). Thus, we examined whether SCD inhibition promoted the expression of ER stress-related genes. The expression of activating transcription factor (*Atf*) 4 and *Atf3* were increased in CD4^+^ T cells following SCD1 inhibition (Fig. 4C). Furthermore, the expression of multiple ER stress-related genes including *Atf4*, *Atf3*, *Perk* and *sXbp1* were down-regulated in CD4^+^ T cells transduced with SCD1-expressing retroviruses (Fig. 5D). Thus, these results suggest that SCD promote T_FH_ cell survival via the regulation of ER stress genes.

**Figure 5.**
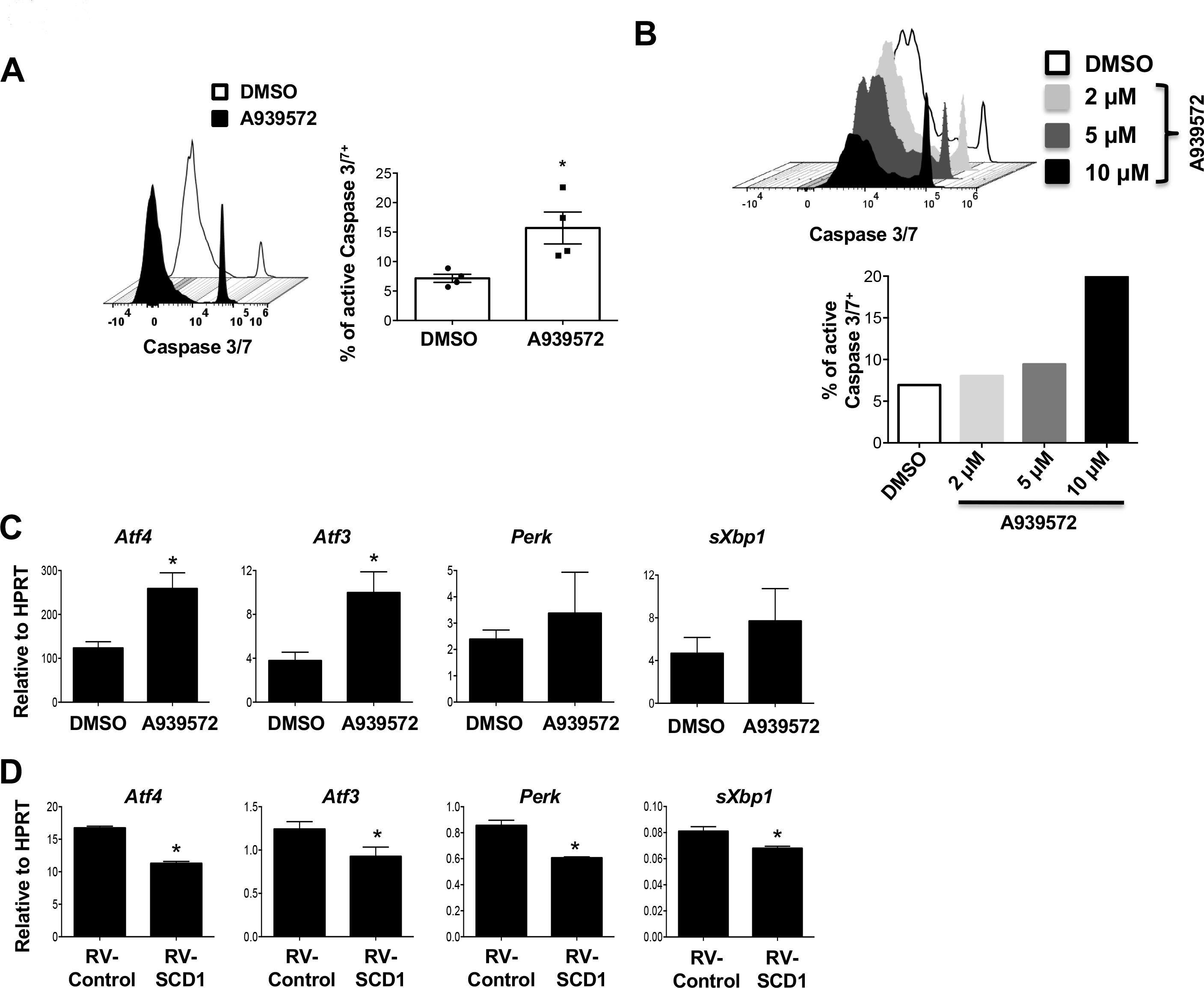
SCD inhibition promotes T_FH_ apoptosis and ER stress gene expression. (A) Caspase 3/7 activity in splenic T_FH_ cells from the mice immunized with X31 virus and treated with SCD inhibitor at 14 d.p.i. (n = 4 - 10). (B) Naïve CD4^+^ T cells were stimulated under T_FH_ polarization condition and treated with increased dosages of SCD inhibitor, active Caspase 3/7^+^ cells were measured at day 4 post activation. (C) The expression of ER-stress related genes in CD4^+^ T cells under T_FH_ polarization with or without SCD inhibitor (10 μM) was determined by qRT-PCR. (D) The expression of ER-stress related genes in CD4^+^ T cells under T_FH_ polarization with or without SCD1 transduction was determined by qRT-PCR. Representative or combine results are obtained from 2 - 3 independent experiments. \**P*<0.05; \*\**P*<0.01; and \*\*\**P*<0.001.

### Enhanced Scd1 expression promotes T_FH_ responses *in vivo*

To identify whether SCD expression contributes cell-intrinsic maintenance of T_FH_ cells, we transduced WT OTII cells with control or SCD1-overexpressing retroviruses. We then adoptively transferred the OT-II cells into CD45.1 congenic mice and infected the mice with X31-OTII virus (Fig. 6A). We found that there were higher levels of T_FH_ cells in SCD1 transduced OTII cells compared to those of control virus transduced cells (Fig. 5B). These data indicate that SCD1 overexpression in OTII cells promotes T_FH_ cell in a cell intrinsic manner.

**Figure 6.**
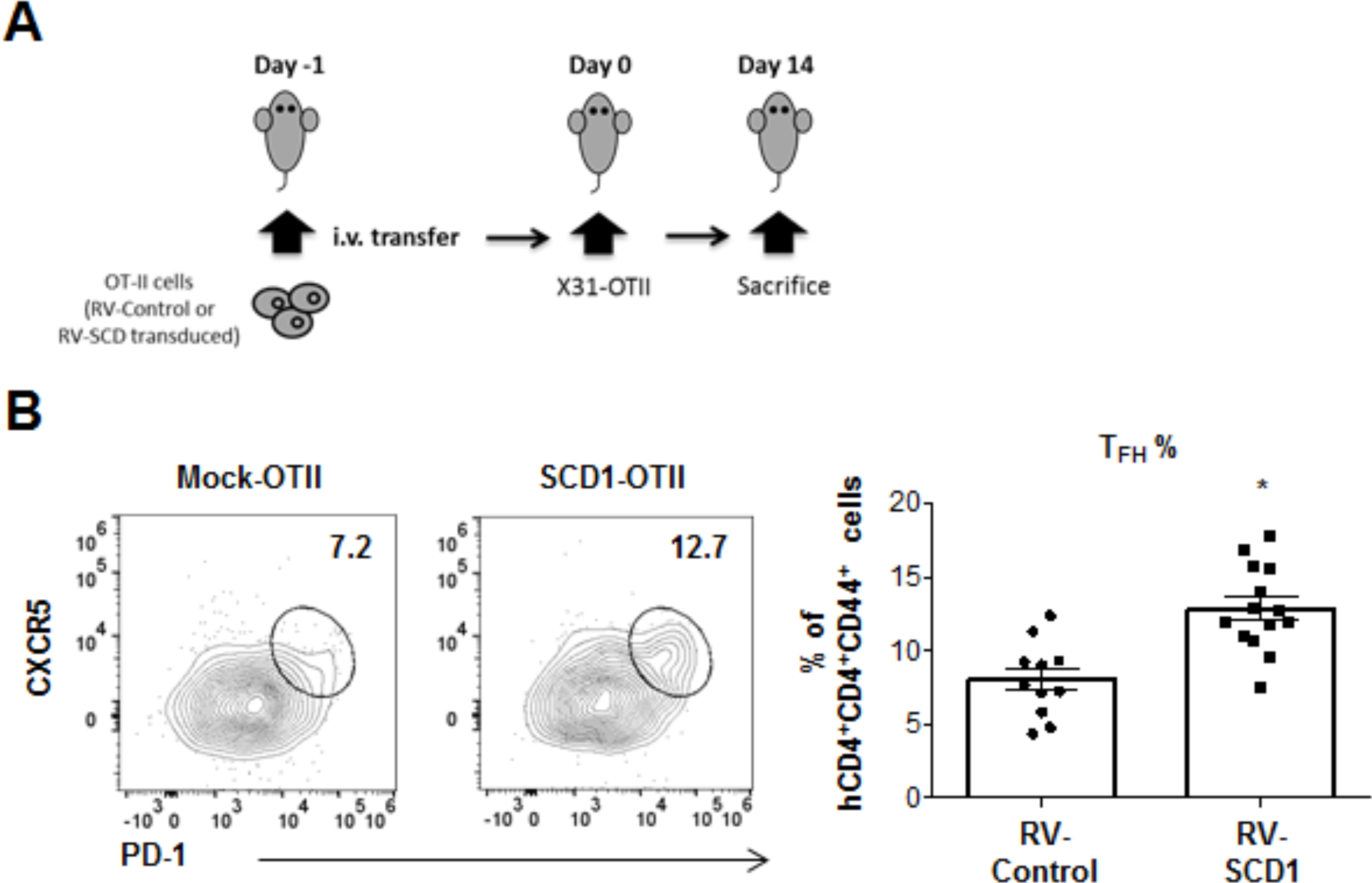
Enforced SCD1 expression in CD4^+^ T cells promotes T_FH_ responses in vivo. Mock or SCD1-retroviral transduced OTII cells were adoptive transferred into CD45.1 congenic mice. Then the mice were immunized with X31-OTII. (A). Schematics of experimental design. (B). The frequency of T_FH_ cells in CD45.2^+^ donor cells were measured in spleen 14 d.p.i.. Results were pooled from 3 independent experiments. * indicates significant differences (*P*<0.05) between control and SCD1-retroviral transduced cells.

## Discussion

In this report, we show that human and murine T_FH_ cells exhibit higher levels of the lipogenic gene expression in the form of *scd1*. The expression of *Scd1* is regulated in a BCL6 dependent manner both *in vivo* and *in vitro* condition. Importantly, a SCD inhibitor, A939572, diminished the magnitude of T_FH_ and GC B cell responses whereas it promoted the frequency of T_FR_ cells *in vivo* following immunization with X31-influenza virus. Therefore, our results have revealed a non-redundant role of SCD and lipid desaturation in regulating T_FH_ maintenance and GC B cell responses.

Excessive or prolonged ER stress, also known as unfolded protein response (UPR), suppresses normal cell growth through inhibition of translation and ultimately apoptosis. Lipotoxicity, induced by overload of lipids and their metabolites, disturbed ER homeostasis and caused metabolic abnormalities and cell death (23). Knockdown of SCD1 enhanced the accumulation of SFAs, resulted in increased expression of ER stress-related genes such as C/EBP homologous protein (CHOP), glucose-regulated protein 78 (GRP78) and splicing of Xbox-binding protein 1 (sXBP1) (22). However, these enhanced ER stress-induced genes and cell death were rescued by treatment of unsaturated fatty acid in Hela cells (24). Interestingly, oleate and palmitoleate, the major products of SCD1 activity, have different metabolic functions. The oleate, but not palmitoleate, normalized the excess ER stress in SCD1 liver-specific deficient mice (11). Thus, SCD expression in T_FH_ cells may contribute cellular homeostasis through maintenance of balance between SFAs and MUFAs. SCD1 have been reported to be highly expressed in several human cancer cells including breast, prostate, lung and ovary (25). SCD inhibitor decreased the proliferation and increased apoptosis in clear cell renal cell carcinoma (ccRCC) *in vivo* and *in vitro* (26). SCD1 inhibition promoted autophagy-induced apoptosis accompanied with autophagy formation in the human hepatocellular carcinoma (HCC) cells via AMPK activation (27). Thus, SCD1 inhibition might be utilized to target cell therapy such as SCD1 high expressing cells including malignant and T_FH_ cells.

Enhanced T_FH_ cells have been reported to be associated with the development and progression of many autoimmune diseases (28). Recently, as the potential target of autoimmune disease therapy, inhibition of lipid metabolism gene has been reported in human patients and animal models. Inhibition of Fasn, one of the fatty acid synthase, reduces development of Th17 resulting in diminished experimental autoimmune encephalomyelitis (EAE) disease progression (29). Simvastatin, inhibitor of HMG-CoA reductase, reduces the serum TNF-α and CRP levels in RA patients (30). In our model, compared to C75 (fasn inhibitor) and fluvastatin, SCD inhibitor (A939572) exhibited superior activities in terms of the inhibition of T_FH_ and GC B cell responsess. Interestingly, it has been reported that obese humans have higher levels of SCD1 expression and MUFA contents compared to lean objects (31). SCD1 has been associated with obesity-related abnormal metabolic diseases such as hyperlipidemia, nonalcoholic liver disease and type 2 diabetes in animal model and human exhibiting increased MUFA levels (11). These reports suggest that T cell lipid metabolism, particularly SCD pathway, may represent a promising target to dampen autoimmunity, especially in those obesity associated autoimmunity (32).

In this study, we show that inhibition of SCD reduces the number of T_FH_ and GC B cells post influenza virus immunization. It is possible that SCD inhibitor may have off-target effects *in vivo* to attenuate T_FH_ and/or GC B cell responses. Furthermore, SCD inhibitor may work through other cell types such as dendritic cells and/or B cells to diminish T_FH_ maintenance. Future studies employing cell-type specific deletion of SCD in different cell types (such as T cells, dendritic cells and/or B cells) are warranted to address these potential limitations of our studies. However, as shown in figure 6, Scd1-overexpression promoted T_FH_ responses *in vivo*, indicating that the promotion of the T cell-intrinsic SCD1 expression facilitates T_FH_ responses *in vivo*.

In conclusion, we found SCD activity promotes T_FH_ cell maintenance by inhibiting their apoptosis and sustaining GC B cell responses. Therefore, SCD may be considered as a novel target to modulate excessive T_FH_ responses during vaccination and/or autoimmunity.

## Acknowledgments

We thank Mayo flow cytometry core for help of cell sorting. This work was supported by the US National Institutes of Health grants (RO1 HL126647, AG047156 and AI112844 to J.S.; RO1 AI132771 to A.D.; T32 AG049672 to N.P.G.) and Kogod Aging Center High Risk Pilot Grant to J.S.

## Additional Information

The authors declare no competing interest.

## Materials and Methods

### Mouse and immunization

Mouse studies were approved by the Institutional Animal Care and Use Committee at the Mayo Clinic or Indiana University School of Medicine (IUSM). All methods were performed in accordance with the relevant guidelines and regulations. WT C57/BL6, CD45.1 congenic, OTII and CD4-cre mice were originally purchased from the Jackson Laboratory and bred in house. BCL6 KO mice (CD4-cre Bcl6^fl/fl^) were generated by crossing CD4-cre transgenic mice with Bcl6^fl/fl^ mice. Bcl6 KO OTII mice were generated by cross OTII transgenic mice to Bcl6 KO mice. Influenza immunization was achieved via intraperitoneal route (I.P.). with Influenza H3N2 A/X31 or X31-OTII (~1.2 × 10^5^ pfu/mouse). A939572 (6 mg/kg, BioFine International, WA, USA), C75 (15 mg/kg, Cayman chemical, MI, USA) or of Fluvastatin (20mg/kg, Cayman chemical) was injected to the X31 immunized mice from day 4 to day 13 for daily via intraperitoneal route.

### Flow cytometry

Splenic T_FH_ and T_FR_ cell were stained with antibodies against mouse CD4 (Clone: RM4-5, Biolegend, San Diego, CA, USA), CD44 (Clone:IM7, Biolegend), PD-1 (Clone: 29F.1A12, Biolegend), CXCR5 conjugated with biotin (Clone: 2G8, BD Biosciences), streptavidin conjugated with APC (Biolegend), Foxp3 (Clone: FJK-16s, Thermo Fisher Scientific, USA), BCL6 (Clone: K112-91, BD Biosciences), ICOS (Clone: C398.4A, Biolegend) and GC B cells were stained with antibodies against mouse B220 (Clone: RA3-6B2, Biolegend), GL7 (Clone: GL7, Biolegend) and CD95 (Clone: SA367H8, Biolegend). For BCL6-retroviral transduced cell gating, anti-mouse H2K antibody (Clone: 36-7-5, BD Biosciences) was used and for SCD1-retroviral transduced cell gating anti-human CD4 antibody (Clone: OKT4, Biolegend) was employed. For intracellular staining, cells were re-stimulated with 100 ng/mL of PMA (Sigma-Aldrich, St Louis, Mo, USA), 1 μg/mL of ionomycin (Sigma-Aldrich) and 40 U/mL of hIL-2 in the presence of 1 μg/mL of Golgistop (BD Biosciences) for 4 hours. Then, cells were stained with surface antibodies and fixed/permeabilization buffers (Biolegend) for staining with anti-IFN-γ antibody. To detect cell apoptosis, the CellEvent Caspase-3/7 Green Detection Reagent (Thermo Fisher Scientific) was added to the cells with complete media for 30 min in 37°C and the caspase-3/7 positive cells were measured on an Attune NXT flow cytometer (Life Technologies, Carlsbad, Calif, USA). For T_FH_ cell sorting, CD4^+^ T cells were enriched from spleen with CD4 microbeads (Clone: L3-T4, Miltenyi Biotec, San Diego, CA, USA). The enriched CD4^+^ T cells were stained with antibodies to define T_FH_ cells as described in flow methods below. And the cells were sorted on a FACSAria (BD Biosciences, San Jose, CA, USA).

### *In vitro* T cell culture

Naïve CD4^+^ T cells were isolated from WT (C57BL/6) or BCL6 flox/flox CD4-Cre (BCL6 KO). Cells were cultured with T_FH_ polarizing condition (pre-coated anti-CD3 (1 μg/ml), soluble anti-CD28 (2 μg/ml), anti-IFN-γ (20 μg/ml), anti-IL-2 (10 μg/ml) antibodies, rmIL-21 (20 ng/ml) and rmIL-6 (20 ng/ml)) for 4 days.

### Quantitative RT-PCR

RNA was extracted by RNA miniprep kit (Sigma-Aldrich) according to the manufacturer’s instructions. Reverse transcription was performed using random primers (Invitrogen, Grand Island, NY, USA) and MMLV (Invitrogen). cDNAs were amplified using SYBR Green PCR Master Mix (Applied Biosystems, Grand Island, NY) using quantistudio 3 PCR system (Applied Biosystems). Data were generated with the comparative threshold cycle method by normalizing to hypoxanthine phosphoribosyltransferase (*HPRT*).

### Retroviral Transduction

Naïve CD4^+^ T or OTII cells were pre-activated with pre-coated anti-CD3 (5 μg/ml) and soluble anti-CD28 (2 μg/ml) antibodies for 2 days. Then cells were transduced with bicistronic retroviruses through spin transduction (2,500 rpm, 90 min). After transduction, cells were cultured with T_FH_ polarized condition for an additional 2 days.

### Adaptive transfer

OTII, BCL6 KO-OTII cells which isolated from BCL6 flox/flox CD4-Cre OTII TCR transgenic mice and SCD1-retroviral transduced OTII cells were adaptive transferred into CD45.1 congenic mice. After one day post transfer, A/X31-OVA_II_ (X31-OTII) (~5.0 × 10^5^ pfu/mouse) was immunized for 14 days via intraperitoneal route.

### ELISA

Levels of X31-specific IgG antibody in the blood serum was measured by enzyme-linked immunosorbent assay (ELISA). In brief, 96-well plates (Biolegend) were pre-coated with 100 *µ*l/well of 7×10^4^ PFU of X31 virus in PBS overnight at 4°C. After blocking with PBS containing 3 % of bovine serum albumin (BSA) (Sigma) for 1 hr at room temperature (RT), each serum sample was serially diluted in blocking buffer to each well and incubated for 2 hrs at RT. After 3 times wash with wash buffer (0.05 % of Tween 20 in PBS), HRP-conjugated goat anti-mouse IgG (Promega, Madison, WI, USA) were added into each well. After incubation for 2 hr at room temperature, tetramethylbenzidine (TMB, Thermo Fisher Scientific) was added to develop the color and then the reaction was stopped by adding 2M H2SO4. The absorbance at wavelength 450 nm was measured by an ELISA reader (Molecular Devices, Sunnyvale, CA, USA). Endpoint titers were calculated as the titer of dilution that gave an absorbance value (O.D.) of 0.2.

### Statistical analysis

Graphs were generated by GraphPad Prism software. Statistical significance was evaluated by calculating ρ-values using one-way ANOVA or Student’s t-test. Significance between the groups was judged based on *p* –value < 0.05 (two tailed).

## Author Contributions

Y.S. performed the experiments, analyzed the data and wrote the manuscript, I.S.C. and N.P.G. performed the experiments and edited the manuscript, A.L.D. provided key reagents and edited the manuscript, J.S. supervised the project, analyzed the data and wrote the manuscript.

---

**Supplementary Fig. 1.**
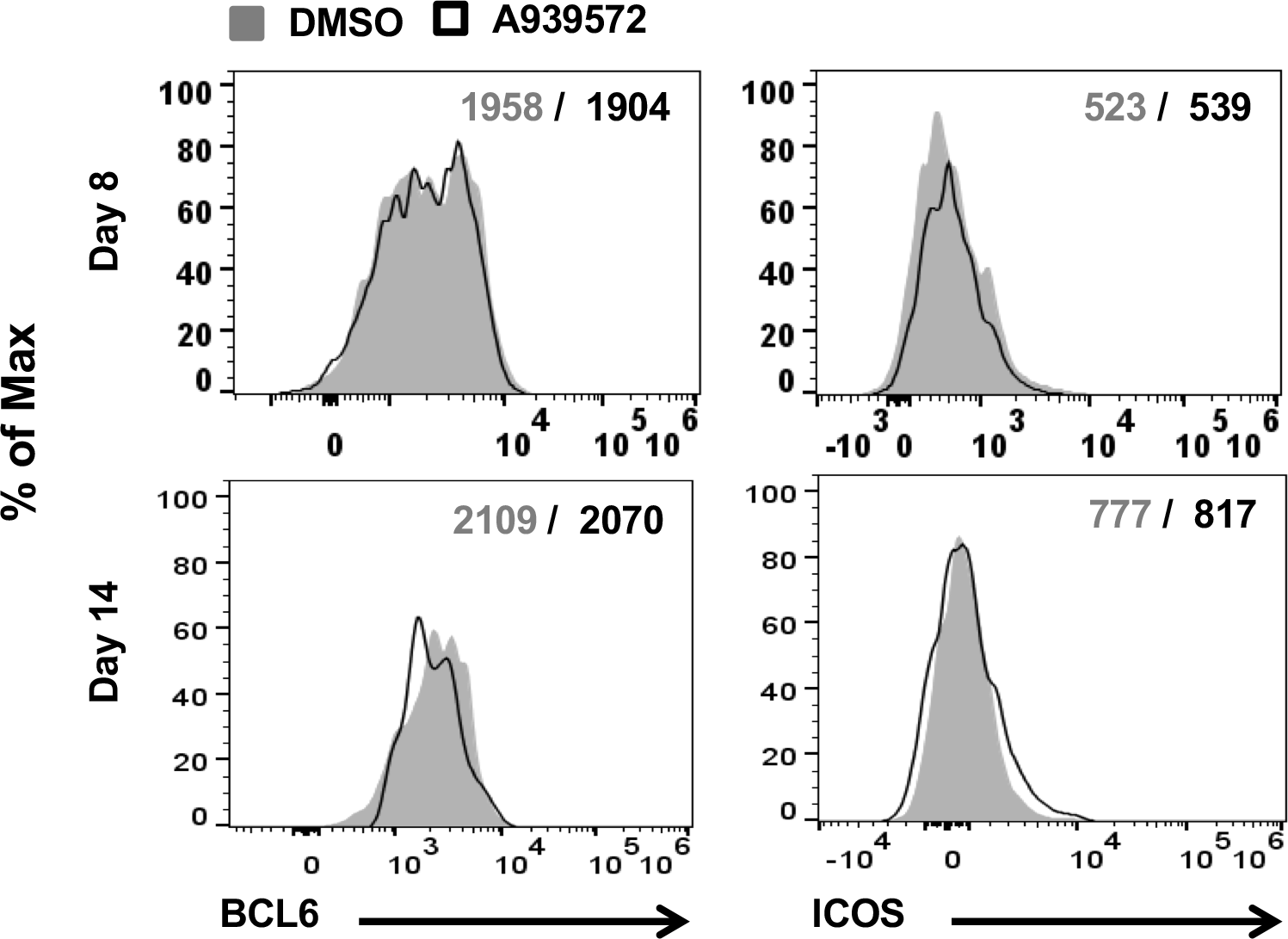

